# Harvestman: A framework for hierarchical feature learning and selection from whole genome sequencing data

**DOI:** 10.1101/2020.03.24.005603

**Authors:** Trevor S. Frisby, Shawn James Baker, Guillaume Marçais, Quang Minh Hoang, Carl Kingsford, Christopher James Langmead

## Abstract

We present Harvestman, a method that takes advantage of hierarchical relationships among the possible biological interpretations and representations of genomic variants to perform automatic feature learning, feature selection, and model building. We demonstrate that Harvestman scales to thousands of genomes comprising more than 84 million variants by processing phase 3 data from the 1000 Genomes Project, the largest publicly available collection of whole genome sequences. Next, using breast cancer data from The Cancer Genome Atlas, we show that Harvestman selects a rich combination of representations that are adapted to the learning task, and performs better than a binary representation of SNPs alone. Finally, we compare Harvestman to existing feature selection methods and demonstrate that our method selects smaller and less redundant feature subsets, while maintaining accuracy of the resulting classifier. The data used is available through either the 1000 Genomes Project or The Cancer Genome Atlas. Access to TCGA data requires the completion of a Data Access Request through the Database of Genotypes and Phenotypes (dbGaP). Binary releases of Harvestman compatible with Linux, Windows, and Mac are available for download at https://github.com/cmlh-gp/Harvestman-public/releases

## 1 Introduction

Supervised learning from high-throughput sequencing data presents many challenges (D’Argenio, 2018; Larrañaga et al., 2006; Leung, Delong, Alipanahi, & Frey, 2016). First among these is the curse of dimensionality, which predisposes learning algorithms to overfitting and imposes barriers to scalability (Clarke et al., 2008). A second critical challenge is that raw variant calls may not be the optimal feature encoding for a given learning task (Libbrecht & Noble, 2015). The most informative, and biologically relevant encoding of a given variant may be at a higher level of organization, such as a perturbation in a particular exon, transcript, or pathway. This paper addresses both challenges by introducing Harvestman, a method that automatically identifies a non-redundant set of relevant features chosen from a hierarchy of biological encodings (i.e., interpretations) of the raw variants.

Strategies for finding effective representations of the data include feature engineering methods, which apply domain knowledge to define features *a priori*, and feature learning methods, which apply supervised or unsupervised learning algorithms to the task. Feature engineering is largely a manual process, but for that reason it is likely to produce encodings that are meaningful to domain experts. Feature learning methods are largely automated, but may produce features that are difficult to understand (Bengio, Courville, & Vincent, 2012; Domingos, 2012). Harvestman employs a hybrid approach to finding an effective encoding of the data. It first constructs a hierarchy of potential representations for each variant. We refer to that hierarchy as the *knowledge graph*. Our knowledge graph is derived from existing genomic annotations and ontologies, to ensure that each putative encoding is biologically relevant, but the Harvestman framework can also incorporate alternative, user-defined knowledge graphs.

Traditional approaches to mitigating the curse of dimensionality fall into two basic categories: feature extraction methods and feature selection methods (Li et al., 2017). Feature extraction methods find a new lower-dimensional feature subspace using information about the current feature space, usually in an unsupervised fashion. Hence, the extracted features are not present within the original data, but rather constructed from it. Among the most common feature extraction techniques is Principle Components Analysis. A down-side with such feature extraction methods is that they may become unreliable when the vast majority of the input features are irrelevant to the prediction task, as is often the case with genomic data (Xing, Jordan, & Karp, 2001). Moreover, the induced features are typically linear or non-linear combinations of the input covariates. Such representations can be difficult to understand.

Feature selection methods, in contrast, explicitly select informative subsets of covariates (Blum & Langley, 1997), and do not change the way those features are encoded. Thus, if the given features are have an intuitive interpretation, the chosen subset will be as well. Feature selection techniques are typically categorized as being either a filter, a wrapper, or an embedded method (Hira & Gillies, 2015; Saeys, Inza, & Larrañaga, 2007). Filter-based methods (e.g., ReliefF (Kononenko, Šimec, & Robnik-Šikonja, 1997)) rank individual features according to some scalar quantity, such as the mutual information between the feature and the label. The user then selects the top *k* features for subsequent model-building. Wrapper methods (e.g., CFS Hall (1999)) explicitly rank various subsets of features, with the highest ranking subset then chosen. Embedded methods select feature subsets during model building (e.g., by applying *L*1-regularization).

Harvestman employs supervised hierarchical feature selection under a wrapper-based regime, as it solves an optimization problem over the knowledge graph designed to select a small and non-redundant subset of maximally informative features. In this way, Harvestman automatically learns the best feature encoding while performing feature selection.

## 2 Background and Related Work

Traditional feature selection strategies are not intended for hierarchical feature spaces, where the parent of a given feature represents an alternative encoding of the same underlying observation(s). Harvestman builds on recent techniques (Ristoski & Paulheim, 2014; Wan & Freitas, 2015; Wang et al., 2017) for solving the hierarchical feature selection problem. Let *V* = {*v*_1_, …, *v*_*m*_} be a set of features (nodes) and let *G* = (*V, E*) be a directed acyclic graph over those features. Here, the topology of *G* encodes the hierarchical relationships among the features (if any) such that a directed edge from node *v*_*i*_ to *v*_*j*_ implies that *v*_*j*_ represents a higher level abstraction of node *v*_*i*_. In the current paper, *G* is the knowledge graph that encodes the potential interpretations (i.e. feature encodings) for a given set of variant calls (see Fig. 1). Naturally, each vertex/feature in *G* will be correlated with its ancestors and descendants, and so it is important to identify and eliminate redundant features. The (supervised) hierarchical feature selection problem is: given *G* and a set of labeled training instances, select the most informative and least redundant subset of *V*.

**Figure 1:**
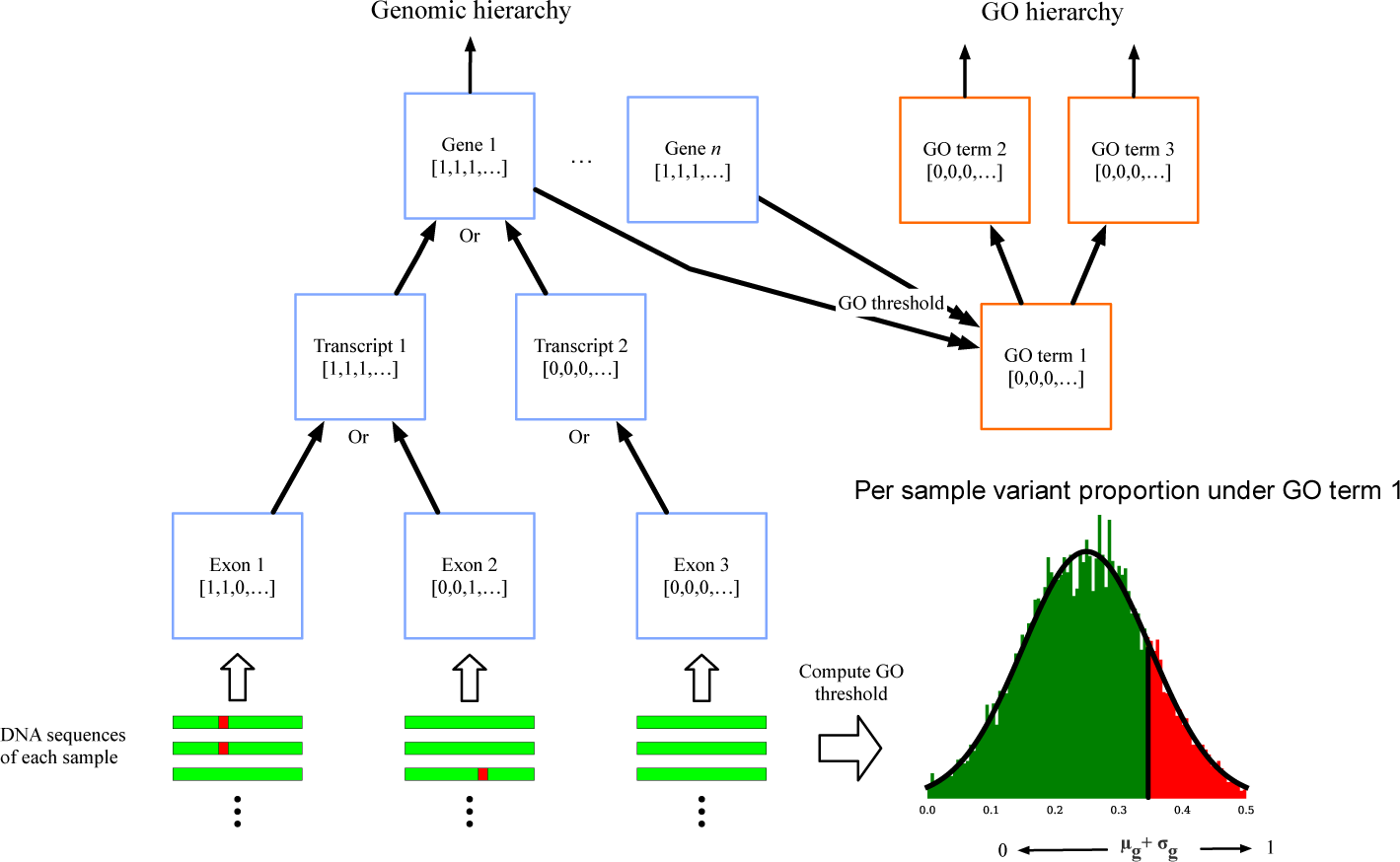
Harvestman**’s knowledge graph and variant encoding scheme**. The knowledge graph is composed of the genomic hierarchy (blue boxes) and GO hierarchy (orange boxes). Binary vectors at the genomic hierarchy leaf nodes are determined directly from DNA sequences (shown by green bars, variants in sequence shown by red boxes). Binary vectors at parent nodes are computed by taking the logical OR of their child nodes or directly from the DNA sequence. A GO threshold is determined for each GO term from variant sequences related to its connected gene nodes (see section 3.1). We use this threshold to determine a binary vector that reflects whether or not each sample is greater or less than the threshold.

There are many different hierarchies that might be used to construct the knowledge graph from genomic data. Perhaps the most obvious one corresponds to the overlap between genomic loci known to play a functional or regulatory role. Here, the terminal nodes of *G* might correspond to specific positions within the genome. Internal nodes correspond to higher levels of annotation that denote specific regions in the genome, such as transcripts or genes. Alternatively, one might use the existing Gene Ontology (GO) hierarchies (Ashburner et al., 2000; The Gene Ontology Consortium, 2017) to define the knowledge graph. The GO graphs describe the cellular components, molecular functions, and biological processes associated with each gene and its products. Unlike genome annotations that relay structural information about specific genomic regions, GO annotations provide broader information about the systems and processes that changes to these regions may affect. Combining the knowledge contained by multiple annotation types thus captures a fuller picture for genomic variation. Harvestman’s strategy is thus to combine any given graphs into a unified hierarchy that represents a wide range of potential interpretations of the raw variants. In the current paper, Harvestman combines a graph extracted from the genome annotation with the three GO graphs.

Hierarchical feature selection is a relatively new area of research. Harvestman is most closely related to a recent greedy algorithm by Ristoski and Paulheim named SHSEL (Ristoski & Paulheim, 2014), as both approaches seek to maximize feature relevance while reducing feature redundancy. The SHSEL algorithm has two steps. In the first step, SHSEL iteratively processes the graph from the leaves to the root. A node is removed from *G* if it is uncorrelated with the label (i.e. irrelevant), or highly correlated with one of its ancestors (i.e. redundant). In the second step, SHSEL computes the average relevance of the remaining nodes along each path from the root to a leaf. A node is removed if its relevance is below-average on a given path. The SHSEL algorithm is elegant, but it is not guaranteed to output an optimal set of features (i.e., those that are both maximally relevant, and least redundant). Like Harvestman, SHSEL is also a supervised approach. While we focus this work on supervised methods, we recognize that unsupervised methods have also been explored (Wang et al., 2017).

Applications of hierarchical feature selection to biology and medicine have been reported, including the HIP (select Hierarchical Information Preserving features) and MR (select Most Relevant) algorithms by Wan and Freitas (2013, 2015, 2018). Like Harvestman, these approaches use GO to define a hierarchy of binary features, although are intended for the analysis of gene expression data rather than genomic variants, and do not incorporate any other hierarchy types. Additionally, HIP and MR are intended for lazy-learning, where feature selection and model building are performed for each new instance. Harvestman instead identifies features that work well across a cohort of samples.

Genetic algorithms have also been used to perform feature selection over the GO hierarchy. The Genetic Algorithm for Hierarchical Feature Selection (GA-HFS) is a wrapper method that identifies an informative subset by applying mutation and selection operations to a virtual population (da Silva, Plastino, & Freitas, 2018). The fitness function used by GA-HFS confers an advantage to informative, yet non-redundant subsets of features. However, like all genetic algorithms, the optimality of the solution is not guaranteed.

Algorithms applied over biological hierarchical structures other than GO have also been explored. Radovanovic, Vukicevic, Kovacevic, Stiglic, and Obradovic (2015) use the hierarchical ICD-9 (International Classification of Disease) codes to predict 30 day hospital readmission. They propose the Group Hierarchical Feature Compression and Selection Method (GHFCS), a greedy bottom up approach that is similar in nature to SHSEL. The algorithm simply removes child nodes from parent nodes if they have lower average information content. GHFCS has been shown to lead to more interpretable feature subsets, as it reduces the total number of features more than competitor methods. However, their algorithm provides no guarantees of optimality.

Other approaches have defined the hierarchical feature selection problem differently than is used in the current paper. A feature and sample selection algorithm by An et al. (2017) uses brain imaging paired with genomic data to predict Alzheimer’s Disease diagnoses. They devise a semi-supervised approach to address inherent redundancy in the data, and use the term hierarchical feature selection to describe the iterative nature of their process for selecting features, rather than the intrinsic structure among the features. Zhao, Chung, Johnson, Moreno, and Long (2016) introduce a hierarchical group penalty for regularized regression settings to identify novel biomarkers in prostate cancer recurrence. Their penalty is similar in spirit to other approaches in statistics such as the fused or sparse lasso, though these previous methods are not capable of handling hierarchical data. The authors note that their approach is computationally expensive, and that quicker algorithms are needed to extend their approach to higher dimensional settings. Similar penalty terms have also been introduced to identify how genetic variation influences gene expression patterns, such as the tree lasso (Kim & Xing, 2012).

As previously mentioned, one limitation of existing hierarchical feature selection methods is that they provide no guarantees with respect to the optimality of the chosen features. Harvestman, in contrast, formulates the problem as an integer-linear program (ILP) where the user specifies an objective function and an optional set of constraints. The objective function defines the desired tradeoff between some measure of feature relevance (e.g., mutual information) and redundancy (e.g., correlation). The user may also specify suitable linear constraints, such as the maximum number of features to be selected. An ILP solver then returns a subset of maximally informative and minimally redundant subset of features, subject to the constraints, or else reports that no solution exists. Ghalwash, Cao, Stojkovic, and Obradovic (2016) propose a similar ILP-based method for (non-hierarchical) feature selection using expression data. They ultimately relax their ILP to a convex optimization problem, because integer programming is NP-complete. We will demonstrate that when using modern ILP solvers, it is possible to perform hierarchical feature selection over very large knowledge graphs. We note that while Harvestman does make simplifying assumptions with the data prior to solving an ILP, the problem presented to the ILP is solved exactly. When we refer to the optimality of Harvestman, we are referring to the value of the user-specified objective.

## 3 Methods

Harvestman performs automatic feature learning, feature selection, and model building in three steps: (i) hierarchy construction; (ii) optimal hierarchical feature selection; and (iii) model building (see Figure 2). These steps are described in the following subsections.

**Figure 2:**
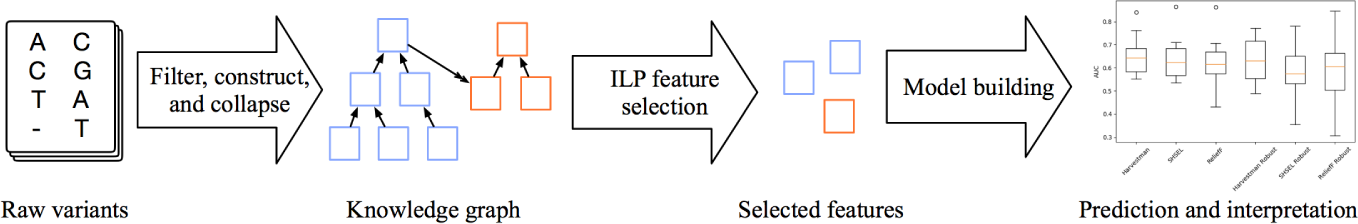
A schematic overview of Harvestman’s analysis pipeline.

### 3.1 Constructing a hierarchy of feature representations

Harvestman leverages hierarchical relationships among potential feature encodings to facilitate feature selection. Figure 1 outlines the process of constructing a knowledge graph from one or more variant call format (VCF) files. The first step encodes annotated variants and structural elements (see Table 1) as a directed acyclic graph. We refer to this initial graph as the ‘genomic hierarchy’. Each node in the genomic hierarchy corresponds to a genomic element. The topology of the graph reflects the logical relationships among these elements. For example, the children of a node representing a gene will be the known transcripts of that gene. Similarly, the children of a node representing a particular transcript will be the exons contained in that transcript.

**Table 1:** The Ensembl IDs that are used by Harvestman, as well as a description of the genomic feature type that they represent.

| Annotation type | Description |
| --- | --- |
| genes | A known protein or RNA coding gene |
| transcripts | A known transcript in a coding gene |
| exons | A known exon in a coding gene |
| peptides | Identifies sequences associated with a known peptide |
| UTR | 5' or 3' untranslated regions |
| bio region | A catch-all annotation describing a genomic region relevant to some biological process |
| SNP | SNPs annotated by NCBI's RefSNP database |

To build the genomic hierarchy, we use Reference SNP (RefSNP) (Wheeler et al., 2007) and Ensembl annotations (Frankish et al., 2017). Ensembl IDs provide unique identifiers that match regions of the genome to structural and functional elements, such as exons, transcripts, genes, and 3’ or 5’ untranslated regions. Inclusion of these elements are motivated by the biological contributions each can make with respect to certain diseases and phenotypes, particularly in the presence of genomic variation.

Variation on the level of exons, transcripts, and genes perhaps have the most clear relationship to observable phenotypes or disease, as each of these elements contribute to the creation of functional proteins from genomic sequence. Alternative splicing could also be explained by variation at this level, and can contribute to cancer (El Marabti & Younis, 2018). 5’ and 3’ untranslated regions are critical regulators of post-transcriptional gene regulation, and genomic variants in these regions are also implicated in cancer (Schuster & Hsieh, 2019). Ensembl also identifies genomic regions with predicted functional roles, including transcription start sites, enhancers, and promoters, or those that are associated with known peptides. These regions are collectively referred to as ‘biological regions’ or ‘peptides’ accordingly, and variation in such sites have also been tied to cancer among other diseases (Farman, Iqbal, Azam, & Saeed, 2018; Hua et al., 2018; Rhie et al., 2018). Associating an Ensembl ID to each node in the graph thus makes clear the biological interpretation of each feature.

Harvestman grafts the three Gene Ontology (GO) hierarchies onto the genomic hierarchy (Fig. 1, orange nodes). GO is a knowledgebase that relates genes to gene products with respect to cellular components, molecular function, and biological process. Since each GO term is associated with a specific set of genes, Harvestman adds a directed edge from each leaf node in the GO graphs to the appropriate ‘gene’ nodes in the genomic hierarchy (i.e. those with annotation ID ‘gene’). These edges create a combined graph that includes the initial genomic hierarchy and the GO hierarchies. We refer to this structure as Harvestman’s knowledge graph.

Harvestman assigns an *n*-element binary vector to each leaf node in the knowledge graph. Here, *n* corresponds to the number of samples within the VCF file(s). For each leaf node, the algorithm sets bit *i* to 1 if the *i*th sample contains a variant associated with that node. Otherwise, the *i*th bit is set to zero. Then, in a bottom-up fashion, the algorithm assigns *n*-element binary vectors to internal nodes by applying a function that combines the vectors from that node’s children. Various functions can be used, including logical `or`’s, `and`’s, `xor`’s, and threshold functions. The framework is general enough that the choice of function can be tailored to suit properties or defining characteristics of specific portions of the graph. If desired, multiple functions can be evaluated in the same knowledge graph through a duplication of parent nodes. For simplicity, when we refer to the knowledge graph in the following sections, we assume each node has been assigned a binary vector.

In our experiments, most vectors were combined by computing a logical `or`. However, a threshold function was used at the nodes that were originally leaves in the GO graphs. Threshold functions were used at these nodes to avoid saturation of the binary vectors (i.e. vectors of all ones). Recall that edges were created between the GO leaf nodes and their associated gene nodes in the genomic hierarchy to create the knowledge graph. Each GO leaf node may be associated with many genes, so it is easy for their binary vectors to become saturated. Saturated vectors are not informative and will therefore never be selected as a feature. If vectors are combined using logical `or`s, then any ancestor of a saturated node will also be saturated and so a single saturated vector may effectively eliminate an entire subgraph from being selected. We avoid this problem by computing a threshold for each GO leaf node. This threshold identifies samples that have an accumulation of genomic variants in a given region of the genome, which which is a hallmark trait of malignancies (Talseth-Palmer & Scott, 2011). A visual example of this procedure is shown in Figure 1. Let *v* be an arbitrary GO leaf node. The threshold associated with *v* is based on the statistics of the number of variants that are observed among the genes connected to *v*. Briefly, let *µ* (resp. *σ*) be the average (resp. standard deviation) of the number of variants from each sample that map to the genes associated with *v*. This is shown as a distribution in Figure 1. We say that sample *i* is enriched in *v* if the number of variants in the *i*th sample that map to any of the genes associated with *v* is ≥ *µ* + *σ*. The *i*th bit of the binary vector associated with *v* is set to one if sample *i* is enriched for *v* (these samples are highlighted in red in Figure 1). Otherwise, it is set to zero (shown in green). This threshold function is only used to compute the binary vectors for the GO leaf nodes. The remaining nodes from the GO hierarchy are assigned binary vectors by computing logical `or`’s.

It is possible for the binary vectors associated with two nodes connected via an edge to be identical. This will occur, for example, if the out-degree of a node is one. When this happens, the two nodes are redundant, and there is no need to consider both of them during feature selection. We handle this situation by “collapsing” redundant nodes (and pathways of redundant nodes) into a single node. This has the benefit of reducing the total number of features that need to be considered during feature selection.

### 3.2 Optimal hierarchical feature selection via Integer Linear Programming

We introduce here an ILP-based approach that identifies relevant and non-redundant features based on the mutual information (MI) between the features and the given label, and the pairwise correlation between features in the knowledge graph.

Let *B*_*i*_ ∈ {0, 1}^*n*^ be the binary vector associated with node *i* in *G*, and let *L* ∈ {0, 1}^*n*^ be a binary label vector encoding, for example, the presence or absence of a particular phenotype. We denote the mutual information (MI) between *B*_*i*_ and *L* as *I*(*B*_*i*_; *L*). This is a measure of feature relevance, as it indicates how much information we gain about the label after observing feature vector *B*_*i*_. More precisely, the MI between two random variables is defined as:

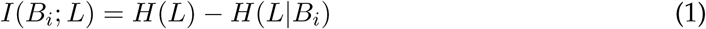

Here, *H*(*L*) is the entropy of the labels, *L*, and *H*(*L*|*B*_*i*_) the conditional entropy of labels, after observing binary feature vector *B*_*i*_. The entropies quantify the uncertainty of the corresponding random variables representing the labels and features. Let *p*(*X*) denote the probability that random variable *X* = 1. Then, for *b* ∈ *B*_*i*_ and *ℓ* ∈ *L* we have the following:

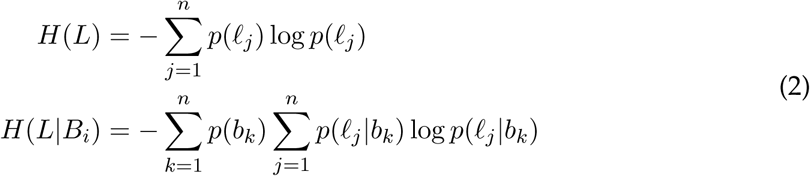

Thus, high MI between feature vector and the label corresponds to lower uncertainty, making such features relevant to classification tasks. MI has commonly and effectively been used as a measure of feature relevance in biological settings (Jansi Rani & Devaraj, 2019; Sun, Zhang, Luo, Cao, & Li, 2019; Zhu, Fan, He, & Xu, 2018).

In practice, we find it useful to pre-filter features using an MI threshold designed to eliminate features that are clearly irrelevant. We consider this an information theoretic equivalent to variant filtration pre-processing steps common to DNA sequence analyses. Harvestman lets the user specify a threshold, *t* ≥ 0, and it pre-filters feature *i* if *I*(*B*_*i*_; *L*) < *t*.

The knowledge graph contains many highly correlated features by construction. Highly correlated features provide little or no additional information and increase model complexity, and so should be eliminated. We denote the correlation between features *i* and *j* as Corr(*B*_*i*_, *B*_*j*_). We used the Pearson correlation coefficient in our experiments, although many other measures of correlation between binary features can be used (Choi, Cha, & Tappert, 2009). Harvestman pre-computes the correlations between all pairs of features that pass the MI filter outlined above.

We include correlations of only a subset 𝒫 of all feature pairs when solving the ILP. We consider correlations between all pairs of features that share a common ‘gene’ node as an ancestor. Additionally, because two different genes may overlap, we also consider correlations between pairs of features that have overlapping gene nodes as ancestors. Next, to account for relationships between nodes across the genomic and GO hierarchies, we include all pairs of features that fall along a directed path within the knowledge graph. Any pair of features fitting the above requirements that are at least moderately correlated (Pearson correlation ≥ 0.3) are included in 𝒫.

The elements of 𝒫 are included as terms in the ILP objective, which is defined as follows:

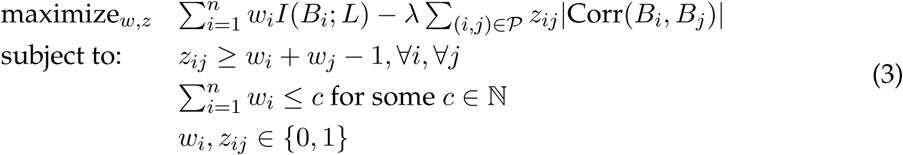

Each feature *i* is associated with a binary decision variable, *w*_*i*_, and each feature pair, (*i, j*), is associated with decision variable *z*_*ij*_. If *w*_*i*_ = 1, then feature *i* is selected. By considering the absolute value of the correlations, we ensure that anti-correlated unselected pairs will not artificially boost the objective value. Parameter λ adjusts the relative importance between mutual information and pairwise correlation. Parameter *c* imposes a constraint on the maximum number of features to select. If *c* is set equal to the number of input features, the ILP will naturally find the optimal number of features to select. In our experiments, we set λ = 1 and varied *c*.

We emphasize that Harvestman’s ILP-based approach to feature selection does not involve the construction and evaluation of classifiers (or any predictive model). Whether a given given feature is selected is determined entirely based on the optimal solution to problem (3).

Finally, Harvestman uses standard Machine Learning libraries to train classifiers and regression models, after the feature selection step. In our experiments, we used Microsoft’s ML.NET machine learning library and scikit-learn (Pedregosa et al., 2011) for model building. We emphasize that model building and evaluation occur after setting aside a hold-out test set. This is done to prevent data leakage from the procedure used to select features to the procedure used to evaluate model performance. To assess model accuracy, we report Area Under the receiver operating Curve (AUC).

## 4 Experiments and Results

We evaluated Harvestman in two ways. First, we tested its scalability using the 1000 Genomes data. Second, we compared Harvestman to existing methods for feature selection on a subset of The Cancer Genome Atlas (TCGA) breast cancer data.

### 4.1 Evaluating Harvestman’s scalability using 2504 whole genome sequences

To demonstrate the effectiveness of Harvestman at scale, we apply our method to data obtained from the 1000 Genomes Project (The 1000 Genomes Project Consortium, 2015), a large and well-known publicly available DNA sequencing data set. In these experiments, we use their most recent Phase 3 data, which includes a combination of low-coverage whole genome and high-coverage exome sequencing for each of 2504 samples. These samples belong to one of five ethnic super-populations (African, American, South Asian, East Asian, and European). In this experiment, we perform feature selection and model building with the task of predicting ethnic super-population from DNA sequence, and evaluate Harvestman’s scalability using a variety of solvers and progressively more powerful public cloud computing instances.

We run five-fold cross validated experiments using two versions of the knowledge graph, one without SNP representation totalling roughly 1.5 million nodes, and one with SNP representation totalling roughly 25 million nodes. In Figure 3 we demonstrate both Harvestman and SHSEL as computationally intensive feature selection methods, and a simple threshold test as a baseline. Memory usage is not shown, since it remains relatively constant across experiments and never exceeds 8 GB. Since the knowledge graph is fully constructed while running the inexpensive threshold test, and training a multi-class logistic regression classifier takes no more than several seconds, we can use threshold timings as a proxy for the time spent loading nodes and edges into memory and computing mutual information.

**Figure 3:**
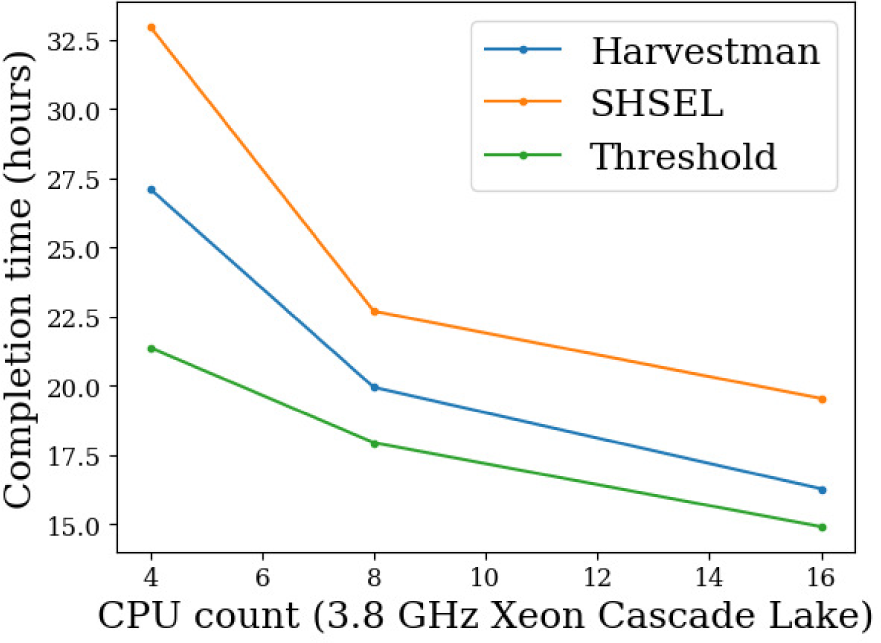
Harvestman **can solve large problem instances in hours** Timing for graph construction and feature selection using Harvestman, SHSEL, and an MI threshold with access to varying numbers of CPUs on the 1000 Genome data. The initial graph consisted of 23,393,068 nodes.

We apply several pre-processing steps before running the feature selection methods. Since building an ILP problem with the full graph would take at least two days of processing time, we filter nodes from the smaller and larger graphs by applying a mutual information threshold of 0.2 and 0.4, respectively. This reduces total feature size to between 15,000 and 30,000 and produces no more than 120,000 correlations to consider. For ease of reproduction, we limit the selection set size to 1000 and use COIN-OR solver with a maximum runtime of one hour.

While running Harvestman to construct an ILP with four CPU cores, graph construction takes 33% of the runtime in the small graph but as much as 79% of the runtime in the SNP-enhanced graph. Harvestman then spends its remaining time computing correlations, setting up an ILP problem, and finally running a solver instance. Once the instance is constructed, even difficult problems can be solved quickly and economically with commercial solvers such as CPLEX. To demonstrate this, we select 5,000 features to predict ancestry on the SNP-inclusive 1000 Genomes graph, which corresponds to an ILP problem with roughly 70,000 considered features and over 1.2 million correlations between them. As shown in Table 2, CPLEX will find an optimal solution in under a minute and six gigabytes of RAM. The commercial Gurobi solver and open source COIN-OR solver will not come to a solution on such a problem in under 48 hours, but are suitable for smaller problems.—

**Table 2:** ILP solve time and maximum memory usage with CPLEX.

| CPU Count | Solve Time (s) | Maximum Memory (MB) |
| --- | --- | --- |
| 4 | 36 | 5179 |
| 8 | 37 | 5714 |
| 16 | 39 | 6569 |

Our method, like SHSEL, scales to multiple processors. Each fold of cross-validation can be run in separate threads that receive data from a single producer thread, which allows us to implement coarse grained parallelism with relative ease. We can achieve finer parallelization by dividing the computation of correlations and other graph processing tasks as we build the ILP. For solving the ILP, both CPLEX and Gurobi allow multiple threads and scaling across multiple machines using message passing. The most time consuming steps that must be performed serially are reading the knowledge graph and associated feature vectors from the binary files, and accessing the solver API to construct the problem from the processed graph. However, these actions can be performed concurrently with the other parts of the program, and do not pose a major hindrance with the number of cores available at the range of a high-end desktop or economically-available cloud instance.

Once the features are selected, training a logistic regression (or any other) classifier takes a trivial amount of time, and its memory consumption increases with the number of features. Models built with the selected features were able to properly classify ethnicity with over 99% accuracy. This high level of accuracy is expected, as it is known that there are strong markers for geographically separated populations scattered throughout the genome (The 1000 Genomes Project Consortium, 2015).

### 4.2 Using Harvestman to predict cancer survival outcomes

A difficult yet important problem in cancer genomics is finding markers that are predictive of patient outcomes. Adding to the difficulty is that the available training data may be small, with respect to the number of patients, and/or imbalanced with respect to the relative proportions of each outcome. We demonstrate here the effectiveness of using our hierarchical representation of DNA sequence data in these settings by building models for two binary breast cancer survival outcomes.

Using a curated subset of the TCGA BRCA cancer data, we considered two binary endpoints: predicting five-year survival and five-year disease-free survival. The five-year survival data set contains 136 samples with a 100/36 outcome ratio. The five-year disease-free survival data set contains 120 samples with an 89/31 outcome ratio. Sequencing and survival status data was obtained from Cooper (2019), and all data was initially processed according to the original TCGA specifications (Network, 2012) We report results obtained from ten different permutations of the data. For each permutation, we held out 30% of the data for testing. We did five-fold cross validation on the remaining 70%, and report the cross-validated accuracy on the holdout set. Thus, each permutation has a unique hold-out set and training set. These ten permutations were created to test the robustness of the feature selection and model building steps. Note that the training and holdout sets for each permutation are not identical between endpoints.

For comparison, we also applied the previously described SHSEL hierarchical feature selection algorithm (Ristoski & Paulheim, 2014) and the ReliefF (Kononenko et al., 1997) algorithm to the same data partitions. ReliefF is a well-known, scalable, but non-hierarchical approach to feature selection. Briefly, ReliefF ranks features according to their ability to discriminate between labels. We note that, being a filter method, RELIEFF simply ranks the features. The user then decides how many features to include in the classifier.

The main tunable parameter for SHSEL is a similarity threshold, which is analogous to Harvestman’s mutual information threshold, *t* (see Sec. 3.2). We used SHSEL’s recommended similarity threshold of 0.99 in our experiments. With RELIEFF, we select the top *c* features, where *c* is the number of features selected by Harvestman when given the same train-test split. We did not place a limit on the number of features Harvestman should choose, rather we allow the ILP to simply choose the number of features that maximize the objective. For each experiment, an identical knowledge graph was used as a starting point for each algorithm. To further show robustness of the method, we report classification accuracy obtained with three different classifier types, logistic regression (LR) with no regularization, random forest (RF) using 100 trees, and support vector machine (SVM) with radial basis function kernel. All unspecified classifier parameters were left in their default settings.

### 4.3 Harvestman’s knowledge graph is more informative than a binary encoding of raw SNPs

Harvestman is predicated on the idea that the knowledge graph, a hierarchical representation of prior knowledge over the human genome, may contain more suitable feature encodings than raw SNPs. By construction, the bottom layer of Harvestman’s hierarchy consists of annotated SNPs, and further loci-centric annotation comprise the higher layers. In order to demonstrate the informative value of the knowledge graph with respect to that of SNPs alone, we ran experiments using three segments of the knowledge graph:

1. All node types
2. All node types except SNPs
3. Only SNPs

For both survival endpoints, we initialized knowledge graphs using initial MI thresholds of 0.05, 0.075, 0.1, and 0.125. For cases (1) and (2) above, we then applied Harvestman’s ILP-based feature selection strategy. We trained classifiers on the selected feature subsets as well as with the SNPs from each graph alone, and report the model AUC as a function of feature counts in Figure 4. Each data point corresponds to one of the four initial knowledge graphs, where the initial MI threshold decreases moving from left to right. We do not perform further feature selection on segment 3 (only SNPs). With respect to the five year survival endpoint, the number of nodes in segment 3 is less than the total number of nodes selected by Harvestman on segments 1 and 2. In contrast, the number of segment 3 nodes is more comparable to those selected on segments 1 and 2.

**Figure 4:**
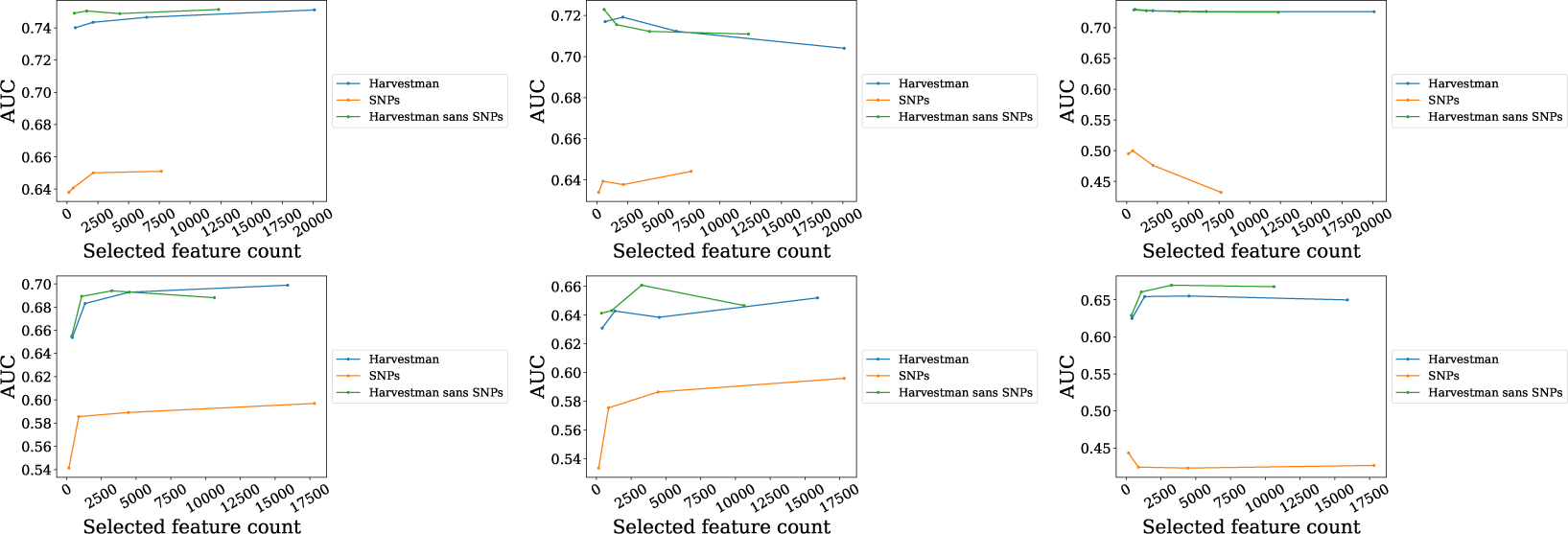
The knowledge graph is more informative than raw SNPs. AUC as a function of feature counts for five year survival (**Top**) and five year disease free survival (**bottom**) obtained with logistic regression (**left**), random forest (**middle**), and SVM (**right**). In each, Harvestman is applied to complete knowledge graph as well as a knowledge graph with SNPs removed using MI thresholds 0.05, 0.075, 0.1, and 0.125. Comparisons are made using models trained on binary encodings of SNPs that passed those same thresholds.

For both endpoints, the features selected by Harvestman from segment 1 and 2 knowledge graphs achieve a higher AUC than those for segment 3 (*p* < 0.05, paired t-tests, see Supplementary Tables S1 and S2), and this behavior generalizes across three classifier types. This suggests that features encoded within the knowledge graph are more informative for these classification tasks than are encodings of SNPs alone. Furthermore, there is no difference in AUC when comparing Harvestman over the complete knowledge graph and Harvestman over the knowledge graph sans SNPs. This further verifies that the best set of features is obtained using portions of the graph representing genomic loci at a broader scale than individual SNPs. If this were not the case, then we would expect Harvestman sans SNPs to perform more poorly than Harvestman over the complete knowledge graph. Additionally, Harvestman is still able to identify these informative, higher level features even when presented less informative SNP nodes. We conclude that the knowledge graph effectively encodes genomic features better than using raw SNPs alone.

### 4.4 Harvestman selects fewer features than SHSEL without sacrificing model AUC

Given the success of using the knowledge graph compared to an encoding of SNPs alone, we next compare Harvestman to SHSEL and RELIEFF over knowledge graphs containing each node type. Using the same initial MI filters as before, we show in Figure 5 the AUCs of three different classifiers on both survival tasks as a function of the number of features selected by each method. We also include as a natural baseline a model trained on all features that passed each initial threshold.

**Figure 5:**
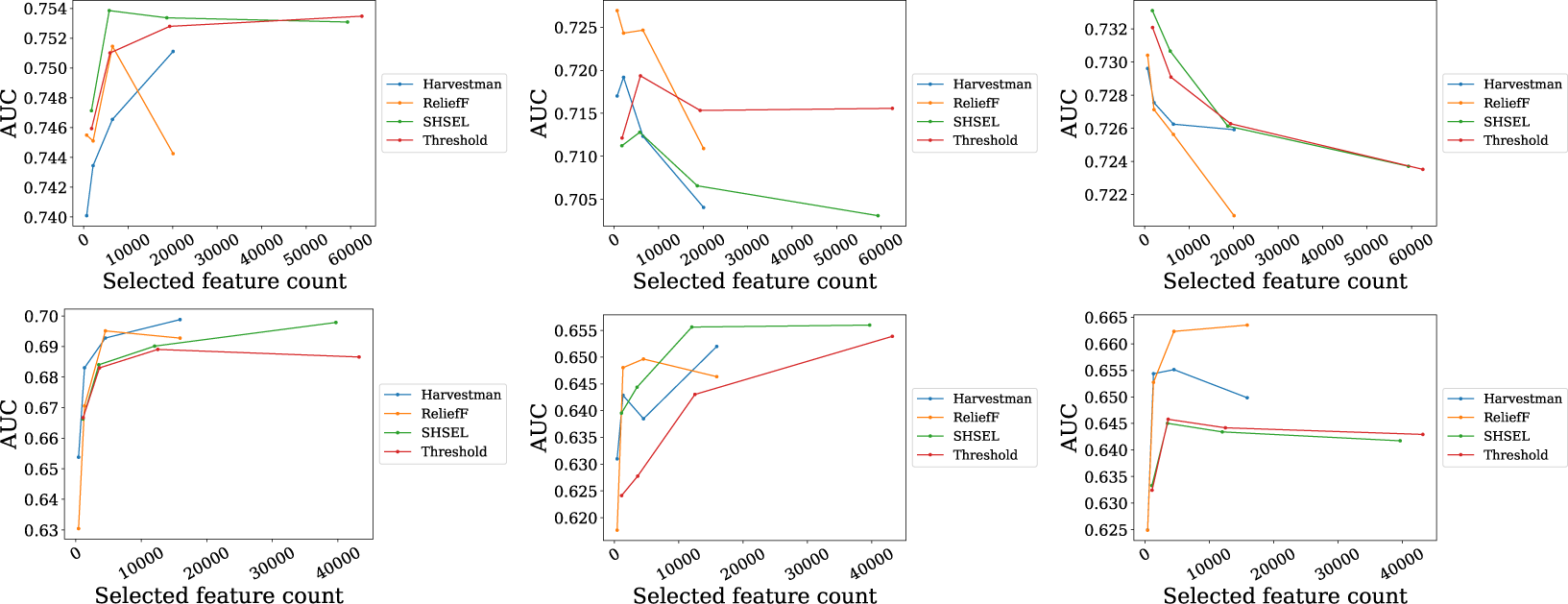
Harvestman **selects fewer features than** SHSEL **without sacrificing model AUC**. AUC as a function of feature counts for five year survival (**Top**) and five year disease free survival (**bottom**) obtained with logistic regression (**left**), random forest (**middle**), and SVM (**right**). In each, Harvestman is applied to complete knowledge graphs with MI thresholds 0.05, 0.075, 0.1, and 0.125.

Reducing the dimensionality of data is the primary goal of any feature selection strategy. In each experiment, we find that Harvestman selects significantly fewer features than SHSEL or the threshold baseline (*p* < 0.05, paired two-sided t-test, see Supplementary Tables S3 and S4). As the initial graph increases in size, this effect becomes increasingly more pronounced. In the case of the most lenient threshold used (0.05), we find that SHSEL selects nearly 60,000 features from the five year survival knowledge graph and 40,000 from the five year disease free survival, and does not improve much upon the threshold baseline. This is compared to about 20,000 and 15,000 features respectively with Harvestman. Thus, Harvestman is more effective, in terms of reducing total feature counts for these two endpoints.

When we consider the underlying AUC of each trained model, we see several patterns that depend on classifier type and the survival endpoint. With respect to disease free survival, there is a general trend that as we decrease the initial threshold, the AUCs tend to increase or plateau, as each feature selection strategy chooses larger numbers a features that are used in training. The highest average AUC measured overall corresponds to Harvestman using LR, though SHSEL and ReliefF obtain the highest AUCs when using RF or SVM, respectively. In most cases, we find no statistically significant differences (*p* < 0.05, paired two-sided t-test, see Supplementary Tables S3 and S4).

In general, we note that all models obtain higher AUC in the five year survival setting compared to five year disease free survival. With the LR classifier, we found no statistically significant differences between model AUCs, though note that the best overall AUC was obtained by a model using SHSEL. RF and SVM classifiers show evidence of overfitting, as AUC tends to decrease as the number of features increases. Note that the scaling of the AUC and feature count axes indicate these differences are not as stark as they may at first seem, as AUC varies by less than 0.2 across experiments using both classifiers. In any case, this trend occurs independent of feature selection strategy. With SVM, there is no statistical difference between AUCs across experiments. With RF, there are instances where a ReliefF model (initial MI threshold 0.125) and Threshold model (initial MI threshold 0.05) outperform the Harvestman model. In the Threshold case, we note that Harvestman had selected significantly fewer features in that experiment. The most common result is that model AUCs obtained with each feature selection method are statistically indistinguishable for a given MI threshold and classifier type. In general, this means that Harvestman can select fewer features than the hierarchical method SHSEL and the baseline threshold method without sacrificing model AUC.

### 4.5 Comparing selected feature sets by average correlation

It is desirable for feature selection algorithms to select non-redundant features. We investigated the redundancy of features selected by each algorithm over knowledge graphs by computing pairwise correlations between subsets of selected features. For each experiment, we randomly selected 1000 features selected by each feature selection strategy, and report in Figure 6 the absolute values of pairwise Pearson correlations between those features. In Supplementary Figures S1 and S2, we show exemplary distributions of these pairwise correlation taken from a single cross-validation experiment from each MI threshold.

**Figure 6:**
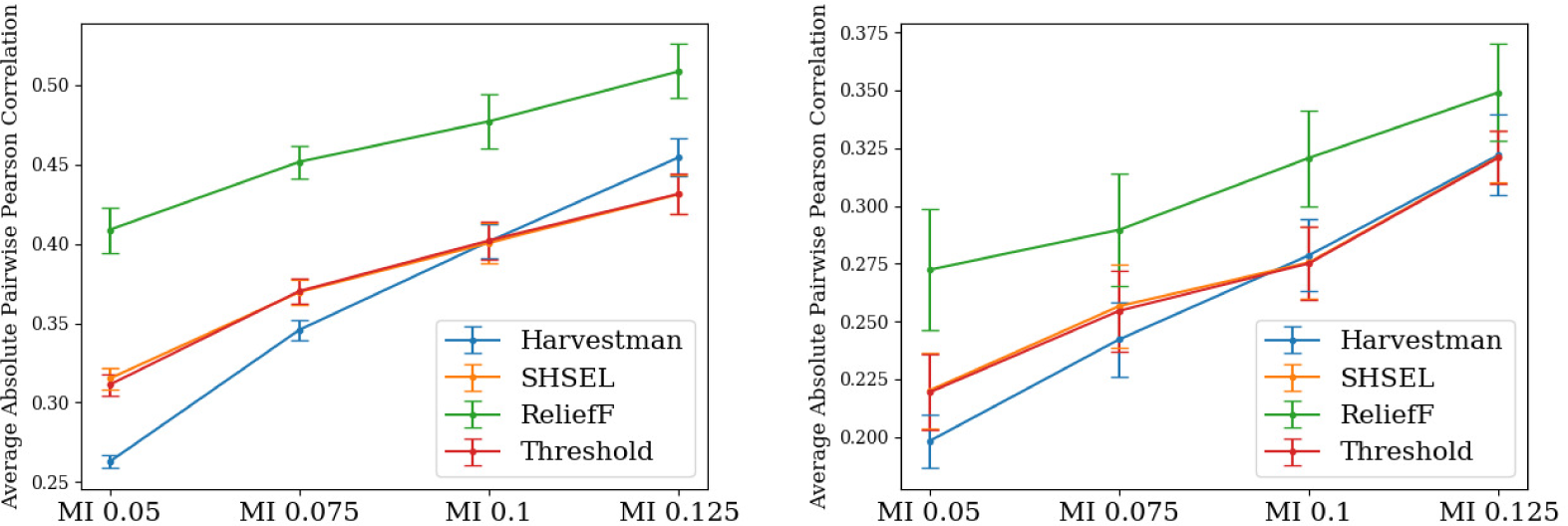
Harvestman **selects fewer redundant features than other methods as graph size increases**. The average absolute pairwise correlation of 1000 randomly sampled selected features for each selection algorithm for both the five year survival (**Left**) and five year disease free survival (**Right**). Error bars denote standard errors over ten runs with different train-test permutations.

In both endpoints, we notice some similar trends. For one, ReliefF consistently selects the most redundant features (Supplementary Tables S5 and S6). While both Harvestman and SHSEL consider pairwise similarity of hierarchically related features as a means of reducing redundancy among their selected feature subsets, ReliefF does not. It therefore makes sense that Harvestman and SHSEL should select less redundant features than ReliefF. Among features that pass the initial mutual information threshold, ReliefF selects a redundant subset, which is why ReliefF also has higher pairwise correlation than the Threshold.

With respect to the two hierarchical feature selection methods, we notice that as the initial MI threshold decreases, there is a more pronounced difference between pairwise correlations obtained by Harvestman relative to SHSEL. For both endpoints, when considering knowledge graphs constructed with MI thresholds 0.05 and 0.075, features selected by Harvestman have lower pairwise correlation compared to SHSEL, and the differences are statistically significant. For thresholds 0.1 and 0.125, Harvestman does not select less redundant subsets. Since lower initial thresholds correspond to larger initial knowledge graphs, this suggests that Harvestman is more adept at finding less redundant features in larger problem instances. Additionally, where Harvestman is able to select fewer redundant features than SHSEL, so too is it able to select fewer features than the MI baseline, whereas SHSEL is unable to improve upon the baseline.

## 5 Discussion and Conclusion

We have introduced Harvestman, a new approach to supervised hierarchical feature selection, and demonstrated it on our knowledge graphs built from high-throughput sequence data. Using the 1000 Genomes, we show Harvestman scales to thousands of genomes. In comparison to alternative methods, Harvestman tends to select smaller or similar sized subsets of features. Smaller feature subsets are desirable for both practical and statistical reasons. In particular, smaller subsets may be easier to understand, and will produce simpler models that are less likely to overfit the training data. Like SHSEL, our method uses a knowledge graph to guide feature selection. However, as the knowledge graph increases in size, Harvestman is more aggressive when it comes to eliminating redundant features.

By construction, each node (feature) in our knowledge graph corresponds to specific genomic annotation. This makes it straightforward to relate prior knowledge to the classification task at hand, and strengthen existing relationships or forge new ones. Moving forward, we look to strengthen our knowledge graph by adding more diverse sets of annotation, particularly those that identify regulatory elements across the genome. This can help make the graph more directly interpretable, and may make inspecting selected features instructive. While our experiments are performed over our knowledge graph, we note that Harvestman and SHSEL are easily configured to use any suitable knowledge graph.

Harvestman provides an additional degree of customization through the ILP objective. In particular, the ILP objective function specifies the global properties of the resulting feature set. For example, users can control the size of the set and adjust the tradeoffs between feature relevance and redundancy. SHSEL, in contrast, performs feature selection via explicit graph traversals. That is, SHSEL selects features based on local properties in the graph, while Harvestman solves a global optimization problem, exactly. Harvestman’s ILP-based formulation also facilitates the enumeration of multiple, distinct solutions, a capability not provided by SHSEL. This means that after the first solution is found, it is possible to force the ILP solver to find a second solution with the same objective value if one exists. In this way, it is possible to generate multiple solutions. In general, it is possible to augment a given ILP objective with arbitrary constraints, and then re-run the solver. In particular, we could add constraints that force the ILP to find a solution that differs from the previous by some minimum number of features. In future work, we envision exploring this functionality as a means to identify robust features by solving iterative ILP problems.

We note that it is straightforward to adapt Harvestman to incorporate additional data types (e.g. expression data). The knowledge graph does not need to be a connected graph. Thus, it is easy to incorporate additional hierarchies, even isolated feature nodes. This same capability means that it is possible to evaluate different ways of constructing internal nodes. In our experiments, the binary vectors associated with the internal nodes of the knowledge graph were primarily constructed using logical `or`’s. If desired, one could create and include additional graphs where the binary vectors are constructed using logical `and`’s or any other user-specified function. This capability may be useful when it is unclear which relationships among features best reflect the true relationships between features. While our experiments were performed on binary-valued features, the approach is capable of incorporating numeric features. In principle, many functions can be used to create parent feature vectors from their child nodes, including those that take and emit real values. As an example, we envision this functionality could be used to create nodes that represent gene expression.

Finally, in evaluating Harvestman’s performance in selecting features and making predictions for survival endpoints in cancer, it is important to consider potential limitations of this data and prediction task. For one, survivorship is a complex issue that has many contributing factors. While genetics certainly plays a role in the likely survival and treatment options available to those with cancer, there are other environmental and lifestyle factors, such as tobacco usage (Ng et al., 2006) or age (Nordenskjöld et al., 2019), that also contribute to survival. Furthermore, the small sample size and imbalanced nature of the data further contribute to making this feature selection and prediction task a difficult enterprise. Still, within this challenging setting, Harvestman was able to identify predictive feature subsets, and thus expose specific genomic markers that may play a role in cancer survival.

## Funding and Acknowledgments

The results here are in part based upon data generated by the TCGA Research Network: https://www.cancer.gov/tcga. This work is supported by the CURE grant 4100070287 from the Pennsylvania Department of Health (PA DOH). The content of this paper is solely the responsibility of the authors and does not necessarily represent the official views of the funding agencies. The Pennsylvania Department of Health specifically disclaims responsibility for any analyses, interpretations or conclusions. This work used the Extreme Science and Engineering Discovery Environment, XSEDE (Towns et al., 2014), which is supported by National Science Foundation grant number ACI-1548562, via the Bridges resource at Pittsburgh Super Computing (PSC) through allocation TG-DMS160012, Big Data for Better Health (BD4BH) in Pennsylvania. This project used the Hillman Cancer Bioinformatics Services, which is supported in part by the National Cancer Institute award P30CA047904. This research is supported by an NIH T32 training grant T32 EB009403 as part of the HHMI-NIBIB Interfaces Initiative. This research is funded in part by the Gordon and Betty Moore Foundation’s Data-Driven Discovery Initiative through Grant GBMF4554 to C.K., and by the US National Institutes of Health (R01GM122935).

## Supplementary Material

**Supplementary Table S1:** P-values from five year survival experiments comparing Harvestman applied to knowledge graphs to binary encodings of SNPs. Paired t-tests over ten train-test data permutations were used. FC refers to feature counts.

|  | MI 0.05 |  |  |  | MI 0.075 |  |  |  | MI 0.1 |  |  |  | MI 0.125 |  |  |  |
| --- | --- | --- | --- | --- | --- | --- | --- | --- | --- | --- | --- | --- | --- | --- | --- | --- |
|  | FC | AUC-LR | AUC-RF | AUC-SVM | FC | AUC-LR | AUC-RF | AUC-SVM | FC | AUC-LR | AUC-RF | AUC-SVM | FC | AUC-LR | AUC-RF | AUC-SVM |
| HARVESTMAN-SNPs | <0.001 | <0.001 | 0.004 | <0.001 | <0.001 | <0.001 | <0.001 | 0.001 | <0.001 | <0.001 | <0.001 | <0.001 | <0.001 | <0.001 | <0.001 | <0.001 |
| HARVESTMAN-HARVESTMAN sans SNPs | <0.001 | 0.972 | 0.239 | 0.694 | <0.001 | 0.304 | 0.99 | 0.739 | 0.002 | 0.283 | 0.554 | 0.944 | 0.012 | 0.221 | 0.333 | 0.583 |

**Supplementary Table S2:** P-values from five year disease free survival experiments comparing Harvestman applied to knowledge graphs to binary encodings of SNPs. Paired t-tests over ten train-test data permutations were used. FC refers to feature counts.

|  | MI 0.05 |  |  |  | MI 0.075 |  |  |  | MI 0.1 |  |  |  | MI 0.125 |  |  |  |
| --- | --- | --- | --- | --- | --- | --- | --- | --- | --- | --- | --- | --- | --- | --- | --- | --- |
|  | FC | AUC-LR | AUC-RF | AUC-SVM | FC | AUC-LR | AUC-RF | AUC-SVM | FC | AUC-LR | AUC-RF | AUC-SVM | FC | AUC-LR | AUC-RF | AUC-SVM |
| HARVESTMAN-SNPs | 0.213 | <0.001 | 0.011 | 0.009 | 0.846 | <0.001 | 0.035 | 0.007 | 0.039 | <0.001 | 0.011 | 0.006 | 0.039 | 0.003 | 0.013 | 0.006 |
| HARVESTMAN-HARVESTMAN sans SNPs | <0.001 | 0.16 | 0.723 | 0.107 | 0.004 | 0.8 | 0.03 | 0.048 | 0.01 | 0.202 | 0.977 | 0.078 | 0.023 | 0.626 | 0.192 | 0.148 |

**Supplementary Table S3:**
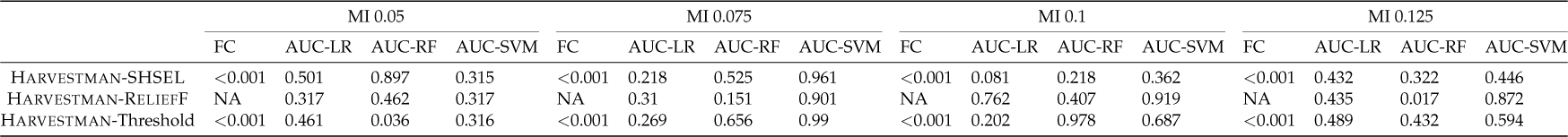
P-values from five year survival experiments comparing Harvestman to SHSEL, ReliefF, and an MI baseline. Paired t-tests over ten train-test data permutations were used. FC refers to feature counts. ReliefF was forced to select the same number of features as Harvestman, hence the NAs.

**Supplementary Table S4:** P-values from five year disease free survival experiments comparing Harvestman to SHSEL, ReliefF, and an MI baseline. Paired t-tests over ten train-test data permutations were used. FC refers to feature counts. ReliefF was forced to select the same number of features as Harvestman, hence the NAs.

|  | MI 0.05 |  |  |  | MI 0.075 |  |  |  | MI 0.1 |  |  |  | MI 0.125 |  |  |  |
| --- | --- | --- | --- | --- | --- | --- | --- | --- | --- | --- | --- | --- | --- | --- | --- | --- |
|  | FC | AUC-LR | AUC-RF | AUC-SVM | FC | AUC-LR | AUC-RF | AUC-SVM | FC | AUC-LR | AUC-RF | AUC-SVM | FC | AUC-LR | AUC-RF | AUC-SVM |
| HARVESTMAN-SHSEL | <0.001 | 0.791 | 0.735 | 0.089 | 0.001 | 0.694 | 0.209 | 0.043 | 0.003 | 0.897 | 0.922 | 0.03 | 0.011 | 0.063 | 0.255 | 0.228 |
| HARVESTMAN-RELIEFF | NA | 0.524 | 0.765 | 0.231 | NA | 0.855 | 0.123 | 0.525 | NA | 0.293 | 0.559 | 0.863 | NA | 0.069 | 0.295 | 0.996 |
| HARVESTMAN-Threshold | <0.001 | 0.204 | 0.896 | 0.095 | 0.001 | 0.48 | 0.661 | 0.042 | 0.004 | 0.996 | 0.103 | 0.022 | 0.012 | 0.096 | 0.394 | 0.342 |

**Supplementary Table S5:** P-values of pairwise correlations of selected features obtained via Harvestman, SHSEL, ReliefF, and an MI threshold for the five year survival endpoint. Paired t-tests over ten train-test data permutations were used.

|  | MI 0.05 | MI 0.075 | MI 0.1 | MI 0.125 |
| --- | --- | --- | --- | --- |
| HARVESTMAN-SHSEL | <0.001 | <0.001 | 0.29 | 0.012 |
| HARVESTMAN-RELIEFF | <0.001 | <0.001 | <0.001 | <0.001 |
| HARVESTMAN-Threshold | <0.001 | <0.001 | 0.866 | 0.144 |
| SHSEL-RELIEFF | <0.001 | <0.001 | <0.001 | <0.001 |
| SHSEL-Threshold | 0.046 | 0.541 | 0.177 | 0.697 |
| Threshold-RELIEFF | <0.001 | <0.001 | <0.001 | <0.001 |

**Supplementary Table S6:**
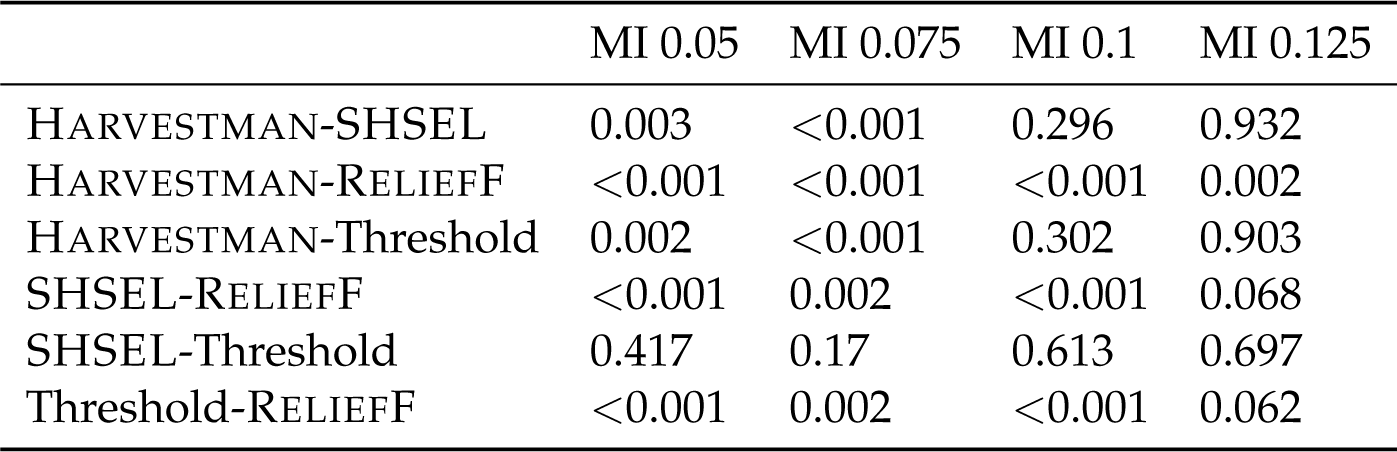
P-values of pairwise correlations of selected features obtained via Harvestman, SHSEL, ReliefF, and an MI threshold for the five year disease free survival endpoint. Paired t-tests over ten train-test data permutations were used.

**Supplementary Figure S1:**
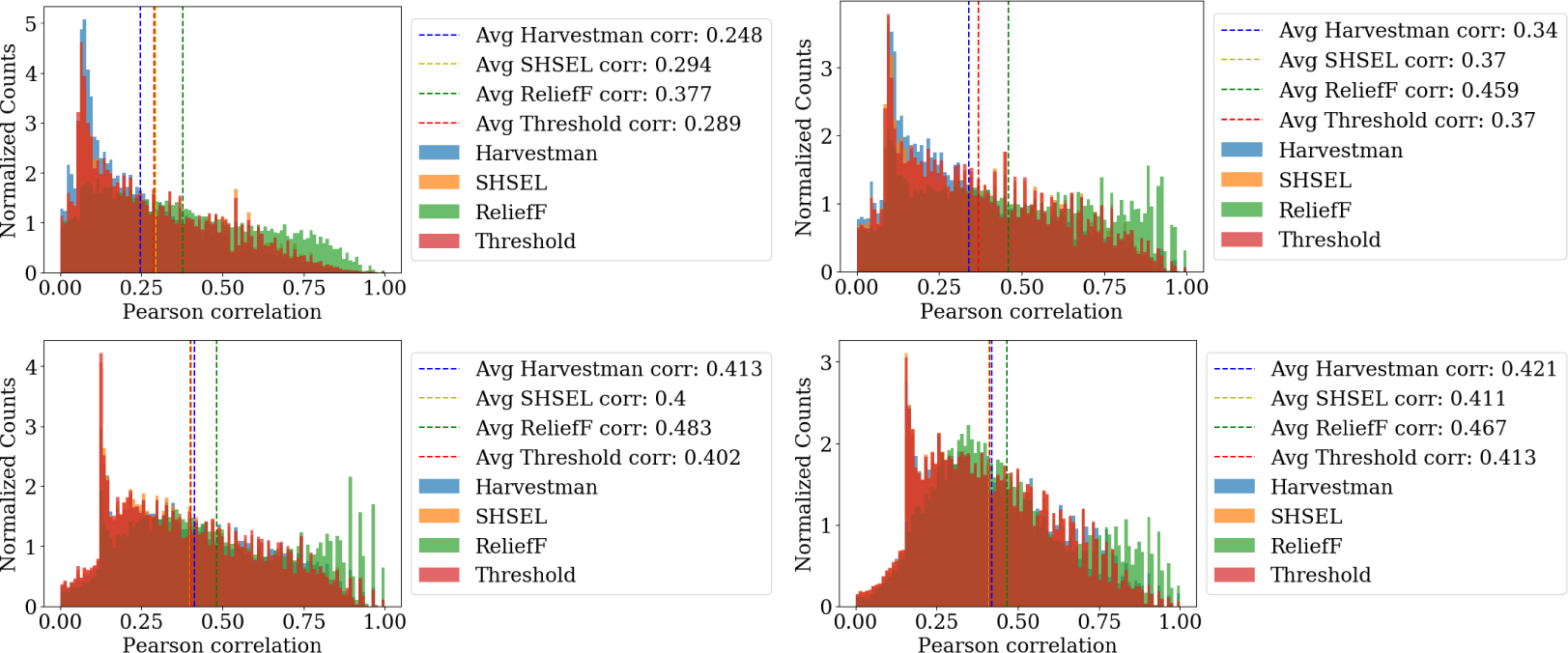
The absolute pairwise Pearson correlations of 1000 random features selected by Harvestman, SHSEL, ReliefF, and an MI threshold from one experiment with the five year survival knowledge graph. The graphs used were obtained from initial MI thresholds of 0.05 (**Top Left**), 0.075 (**Top Right**), 0.1 (**Bottom Left**), and 0.125 (**Bottom Right**).

**Supplementary Figure S2:**
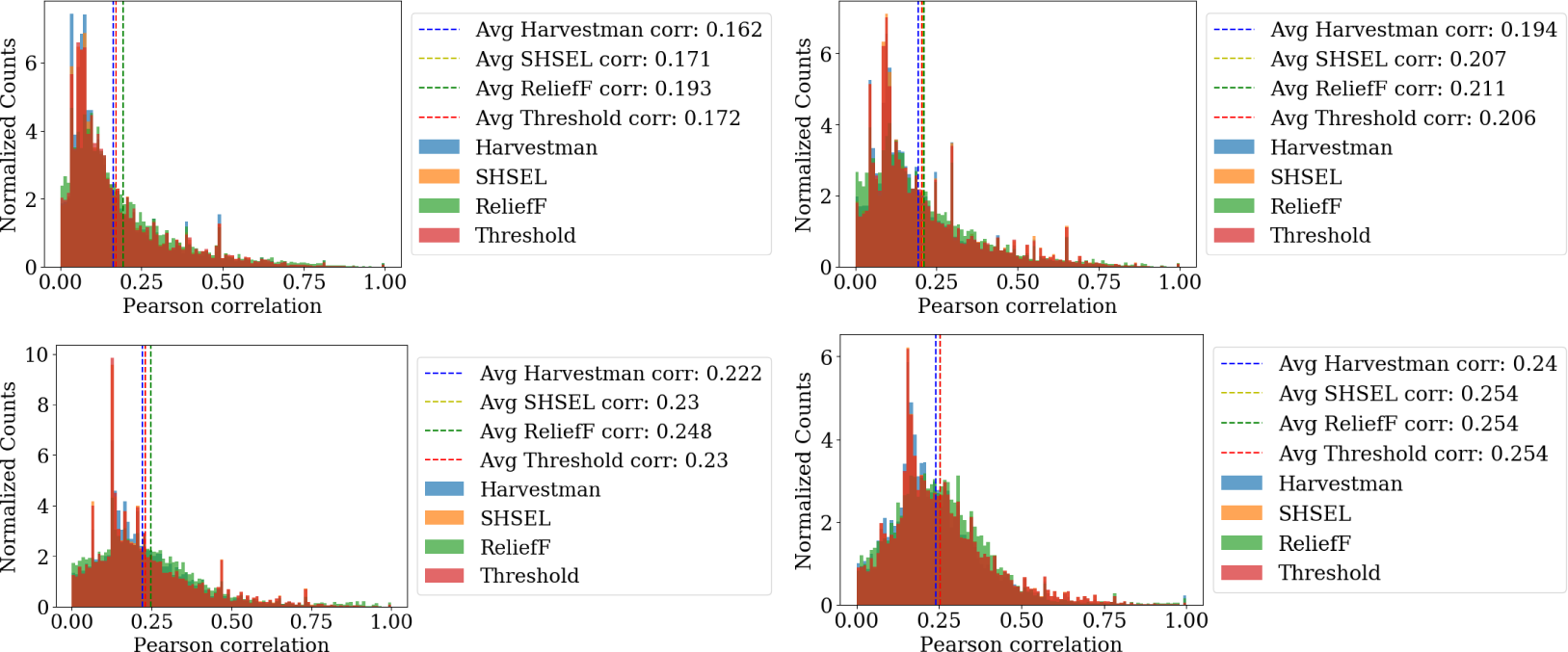
The absolute pairwise Pearson correlations of 1000 random features selected by Harvestman, SHSEL, ReliefF, and an MI threshold from one experiment with the five disease free survival knowledge graph. The graphs used were obtained from initial MI thresholds of 0.05 (**Top Left**), 0.075 (**Top Right**), 0.1 (**Bottom Left**), and 0.125 (**Bottom Right**).

